# Combining tensor decomposition and time warping models for multi-neuronal spike train analysis

**DOI:** 10.1101/2020.03.02.974014

**Authors:** Alex H. Williams

## Abstract

Recordings from large neural populations are becoming an increasingly popular and accessible method in experimental neuroscience. While the activity of individual neurons is often too stochastic to interrogate circuit function on a moment-by-moment basis, multi-neuronal recordings enable us to do so by pooling statistical power across many cells. For example, groups of neurons often exhibit correlated gain or amplitude modulation across trials, which can be statistically formalized in a tensor decomposition framework (Williams et al. 2018). Additionally, the time course of neural population dynamics can be shifted or stretched/compressed, which can be modeled by time warping methods (Williams et al. 2020). Here, I describe how these two modeling frameworks can be combined, and show some evidence that doing so can be highly advantageous for practical neural data analysis—for example, the presence of random time shifts hampers the performance and interpretability of tensor decomposition, while a time-shifted variant of this model corrects for these disruptions and uncovers ground truth structure in simulated data.

## Introduction

Even under highly constrained conditions, animal cognition and behavior is variable over repeated experimental trials. While neuroscientists have historically focused on the trial-averaged firing rates of single neurons, the proliferation of high-density neural recording technologies (T. H. Kim et al. 2016; Stringer et al. 2018; Chen et al. 2018) has fueled research into a variety of interesting single-trial phenomena, such as fluctuations in attentiveness (Cohen and Maunsell 2011), changes-of-mind during decision-making tasks (Kaufman et al. 2015), and incremental learning over the course of many trials (Peters et al. 2014). To understand these, and many other, single-trial phenomena, neuroscientists have cultivated a diverse catalogue of models that pool statistical power across large populations of individually noisy and unreliable neurons (Churchland et al. 2007; Petreska et al. 2011; Pandarinath et al. 2018; Duncker and Sahani 2018; Williams et al. 2018; Williams et al. 2020).

Among the simplest and most common forms of trial-to-trial variability are variability in the amplitude and the latency of a neural response. For example, in the early olfactory system, odor concentration correlates with the amplitude (higher concentrations evoke higher peak firing rates) and latency (higher concentrations evoke more rapid responses) of responses in olfactory sensory neurons and mitral-tufted cells (Olsen et al. 2010; Wilson et al. 2017). Moreover, the access of odorants to olfactory receptors is tightly controlled by airflow through the nose, and thus trial-to-trial variability in the inspiration-exhalation cycle causes additional variability in the latencies of sensory responses (Shusterman et al. 2011; Shusterman et al. 2018). Finally, the amplitude of odor-evoked responses in mitral-tufted cells is subject to experience-dependent modification over short timescales lasting seconds to minutes (often called *adaptation* or *habituation;* Chaudhury et al. 2010) as well as longer-term effects lasting multiple days (Kato et al. 2012). Similar results have been observed in other sensory and motor systems (Salinas and Thier 2000; Dean et al. 2005; Wark et al. 2007; Niell and Stryker 2010; Goris et al. 2014).

In recent work myself and collaborators explored general-purpose statistical models that capture these two complementary forms of single-trial variability (fig. 1). First, in Williams et al. (2018) we studied the possibility of correlated amplitude variability or gain modulation amongst sub-populations of neurons. We found that such a model could be formalized as a well-studied **tensor decomposition** problem (T. Kolda and Bader 2009). Second, in Williams et al. (2020) we studied variability in the timing of dynamics, and formalized this possibility as a **time warping** model. These single-trial variations are statistically detectable when they are correlated across populations of co-recorded neurons.

**Figure 1:**
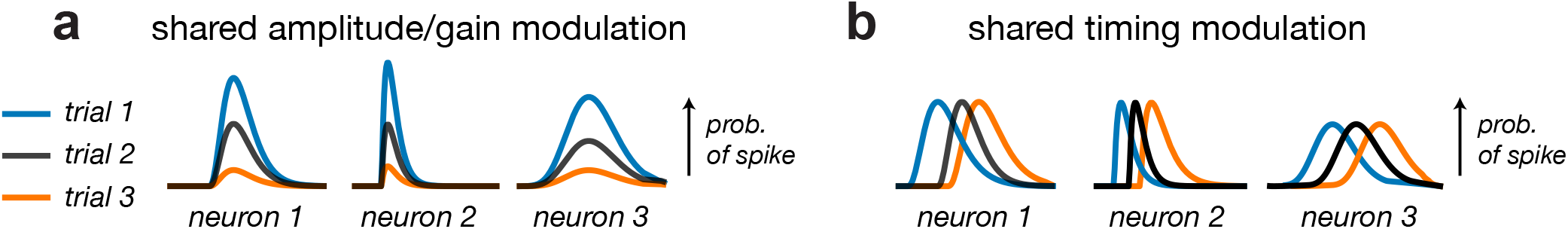
Types of single-trial variation. Colored lines schematize time-varying firing rate functions representing neural responses across multiple trials. (A) Three neurons with correlated amplitudes across trials—i.e., all neurons are either enhanced or suppressed together. Tensor decomposition may be a useful model in cases like this. (B) Three neurons with correlated time delays across trials—i.e., all neurons are either delayed or advanced together. Time warping may be a useful model in cases like this.

We explored the merits of these models in separate papers for clarity, but it is a natural question to ask whether thay can be combined into a unified framework. Indeed, for many experiments we would expect *both* the amplitude and timing of neural dynamics to vary from trial-to-trial. In such cases, neither of the models studied in Williams et al. (2018) and Williams et al. (2020) would provide a satisfactory description of the data. In fact, it is simple to devise scenarios where both models fail to capture any interpretable structure at all, while a unified model that simultaneously accounts for variability in amplitude and timing succeeds.

This manuscript first provides a self-contained summary of tensor decomposition (Williams et al. 2018) and time warping (Williams et al. 2020) models. Two unified models are then outlined: the first extends tensor decomposition by adding per-neuron and per-trial temporal shift parameters (I refer to this as **time-shifted tensor decomposition**); the second incorporates additional dynamical components into the time warping model (I refer to this as a **multi-shift model**).

Previous works have explored similar ideas, though with different motivating applications or proposed algorithms (Harshman et al. 2003; S. Hong and Harshman 2003; Mørup et al. 2008; S. Hong 2009; Q. Wu et al. 2014). In chemometrics, for example, absorption and emission spectra are often shifted due to measurement variability across samples, and this has been incorporated into tensor decomposition models of these datasets (Harshman et al. 2003; S. Hong 2009). In fMRI data, Mørup et al. (2008) introduced and applied a shifted tensor decomposition model with periodic boundary conditions— while periodic boundaries ameliorate computational costs, it may be inappropriate in many cases where evoked neural responses are transient and not rhythmic or periodic. A more recent paper by Duncker and Sahani (2018) incorporates warping functions into a low-dimensional Gaussian Process factor model for spike train analysis; they fit a single nonlinear warping function to all components on each trial, whereas the models I describe apply separate shift parameters to each component, thus enabling the possibility of discovering sub-populations of neurons that are independently time-shifted on a trial-by-trial basis. I also incorporate the possibility that individual units or neurons have characteristic time-shifts in their dynamics, which is not considered by Mørup et al. (2008) or Duncker and Sahani (2018). Additionally, I describe how time warping, tensor decomposition, and hybrids of these two models can be fit by a common optimization strategy based on coordinate and block-coordinate descent (Wright 2015). Overall, this paper aims to synthesize multiple prior works into a unified conceptual framework so that they can be readily understood and applied by practitioners in neuroscience.

## 1 Models

This section provides overview of tensor decomposition (section 1.2; Williams et al. 2018) and time warping (section 1.3; Williams et al. 2020) models of neural activity. Then, two new models of neural activity are introduced—time-shifted tensor decomposition (section 1.4) and multi-shift modeling (section 1.5)—which combine different elements of tensor decomposition and time warping. These two models are closely related to previous works from the field of chemometrics Harshman et al. 2003; S. Hong and Harshman 2003; S. Hong 2009; Q. Wu et al. 2014 and neuroimaging Mørup et al. 2008. Parameter estimation strategies for each of these models are then provided in Section 2.

### 1.1 Notation

Let *X_tnk_* denote the activity of neuron *n*, on trial *k*, in time bin *t*. Let *N* denote the total number of neurons, *K* be the total number of trials, and *T* be the number of timebins in each trial. While we will focus on multi-neuronal recordings as a motivating example, it should be emphasized that the methods described here are applicable to *any* multi-dimensional time series with repeated trials. For example, in the case of behavioral time series, *X_tnk_* could denote the position of body marker *n*, on trial *k*, at time bin *t*. These methods could also be applied to fMRI data with *n* indexing over voxels, *k* indexing trials, and *t* indexing timebins (Mørup et al. 2008).

Lowercase bold symbols denote vectors, e.g. a vector with *n* elements: 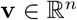. Uppercase bold symbols denote matrices, e.g. a matrix with *m* rows and *n* columns: 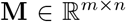. We denote matrix transposes and inverses as **M**^T^ and **M**^-1^, respectively. The trace of a square matrix is denoted Tr[**M**]. We will frequently refer to slices of arrays. For example, 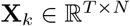 will denote the activity of all neurons on trial *k*, and 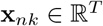 will denote the activity of the *n*^th^ neuron on the *k*^th^ trial. These slicing operations should be familiar to readers that use scientific programming languages; for example, in Python **X**_*k*_ and **x**_*nk*_ respectively correspond the indexing operations `X[:, :, k] and X[:, n, k]`.

Our goal is to define a low-dimensional and interpretable model that approximates this multi-trial dataset. We will use 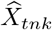 to denote the predicted activity of neuron *n*, on trial *k*, in timebin *t*. For simplicity, will will assume a quadratic loss function (least-squares criterion) throughout this manuscript:

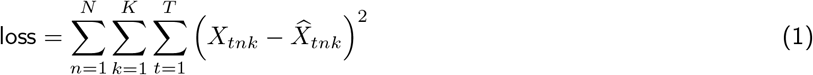

Minimizing this quadratic loss is equivalent to performing maximum likelihood inference in a probabilistic model with Gaussian noise. Different loss functions can be incorporated into tensor decomposition and time warping models (Chi and T. G. Kolda 2012; D. Hong et al. 2018). In particular, a Poisson likelihood criterion is a popular loss function to use for binned spike counts, particularly in low firing rate regimes (Paninski 2004). However, the quadratic loss enables much simpler and faster optimization routines, and I provide some empirical evidence that this loss function can perform well even with low firing rates (see fig. 4; see also Fig 6 in Williams et al. 2020). A more thorough empirical comparison of potential loss functions on spike train data is a worthy direction of future research.

### 1.2 Tensor Decomposition

In Williams et al. (2018) the following **tensor decomposition model** is proposed for neural data:

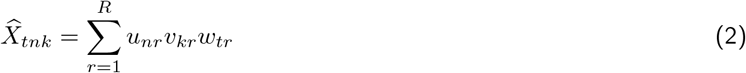

This model approximates the data with *R* components; this is also called a rank-*R* decomposition or a rank-*R* model. The variables *u_nr_* provide a low-dimensional description of the measured features (*neuron factors*); the variables *v_kr_* provide a low-dimensional description of each trial (*trial factors*); the variables *w_tr_* provide a low-dimensional description of the temporal dynamics within every trial (*temporal factors).* This model is schematically illustrated in fig. 2. This model is also known as Canonical Polyadic (CP) tensor decomposition, parallel factor analysis (PARAFAC), and tensor components analysis (TCA). This general method dates back to (Carroll and Chang 1970); see T. Kolda and Bader (2009) for a contemporary review.

**Figure 2:**
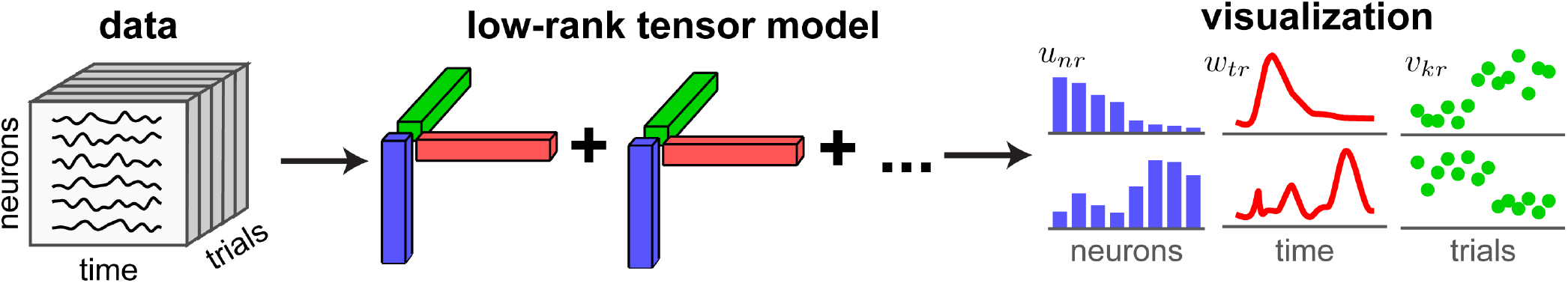
Tensor decomposition model. In applied mathematics, a 3d array of data (left) is often called a third-order tensor. This array can be approximated by a sum of *R* components (middle) which correspond to rank-1 outer products of low-dimensional factors. These factors can be visualized to identify sub-populations of neurons (in blue) that share a prototypical firing pattern within each trial (in red) as well as shared across-trial amplitude modulation (in green). An example with *R* = 2 components is shown on the right.

It is useful to reformulate eq. (2) in terms of standard matrix/vector operations. One can show, for example, that eq. (2) is equivalent to:

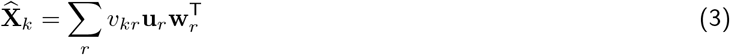

where 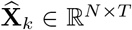 denotes the estimated population activity on trial *k*; 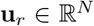 and 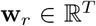 respectively denote the neural factor and temporal factor for the *r*^th^ component.

We will see that eq. (3) is a useful formulation when drawing conceptual connections between tensor decomposition and time warping models. Additionally, this reformulation makes the role of *v_kr_* as a per-trial gain parameter readily apparent— the full population activity on trial *k* is modeled as a linear combination of rank-one matrices, 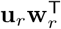, with weights given by *v_kr_*. In the general case, *v_kr_* may be positive or negative, which perhaps complicates our interpretation of these parameters as per-trial “amplitudes,” since one does not typically consider amplitudes to be negative. However, it is often desirable to constrain the factor parameters to be nonnegative anyways (see section 1.2.1), and in this case the interpretation of *v_kr_* as an amplitude is clear. This shared amplitude modulation (also called “gain modulation”) among sub-populations of neurons is consistent with a variety of experimental observations and theoretical models of neural circuits (Salinas and Thier 2000; Goris et al. 2014; Carandini and Heeger 2011; Rabinowitz et al. 2015).

This tensor decomposition model has several other attractive properties. First, it is a relatively straightforward generalization of principal components analysis (PCA), which is already a popular method in neural data analysis. Second, while tensor decomposition is related to PCA, it has favorable properties from the standpoint of *statistical identifiability.* Specifically, under mild conditions the decomposition is *essentially unique*, meaning that non-orthogonal features can be identified by this method (Kruskal 1977; Rhodes 2010; Lim and Comon 2009). In contrast, the reconstruction error of a PCA model is invariant to rotations and invertible linear transformations of the factors, which limits the interpretability of the model. Third and finally, the tensor decomposition model contains exponentially fewer parameters than a PCA model, since it simultaneously reduces the dimensionality of the data across all three axes of the data array (neurons, timepoints, and trials). Thus, tensor decomposition produces a much more aggressive compression of the data, leading to more digestable and visualizable summary of the data. See Williams et al. (2018) for further details and discussion.

In principle, tensor decomposition can be applied to any multi-neuronal recording with a repeated trial structure; however, in certain situations it may not produce a low-dimensional description of the data that is both accurate and interpretable. For example, an accurate tensor decomposition may require a large number of components when (a) neurons within an ensemble fire at slightly different times, or (b) neural ensembles are activated at different times on each trial. Intuitively, these two scenarios result in high-dimensional structure across neurons and high-dimensional structure across trials, respectively. Since tensor decomposition assumes that the data are simultaneously low-dimensional across neurons, trials, and timebins, *high-dimensional structure across any one of these axes may cause difficulties.*

Figure 3 provides a visual illustration of these two “failure modes” of tensor decomposition in a hypothetical dataset containing *N* =10 neurons, *T* =12 timebins, and *K* = 5 trials. First, fig. 3a shows a scenario that is well-modeled by a tensor decomposition with *R* = 1 component. Figure 3b shows the same dataset but with a random delay introduced on each trial, producing high-dimensional structure across trials—unlike fig. 3a, each trial’s activity in fig. 3b is not a re-scaled version of a single rank-1 matrix template. Further complexity is added in fig. 3c, by adding an additional random delay to each neuron’s response. This results in high-dimensional structure across both trials and neurons. However, the multi-trial activity patterns in fig. 3b-c are both intuitively “close” to a rank-1 tensor model. Section 1.4 describes statistical modeling strategies for datasets like this.

**Figure 3:**
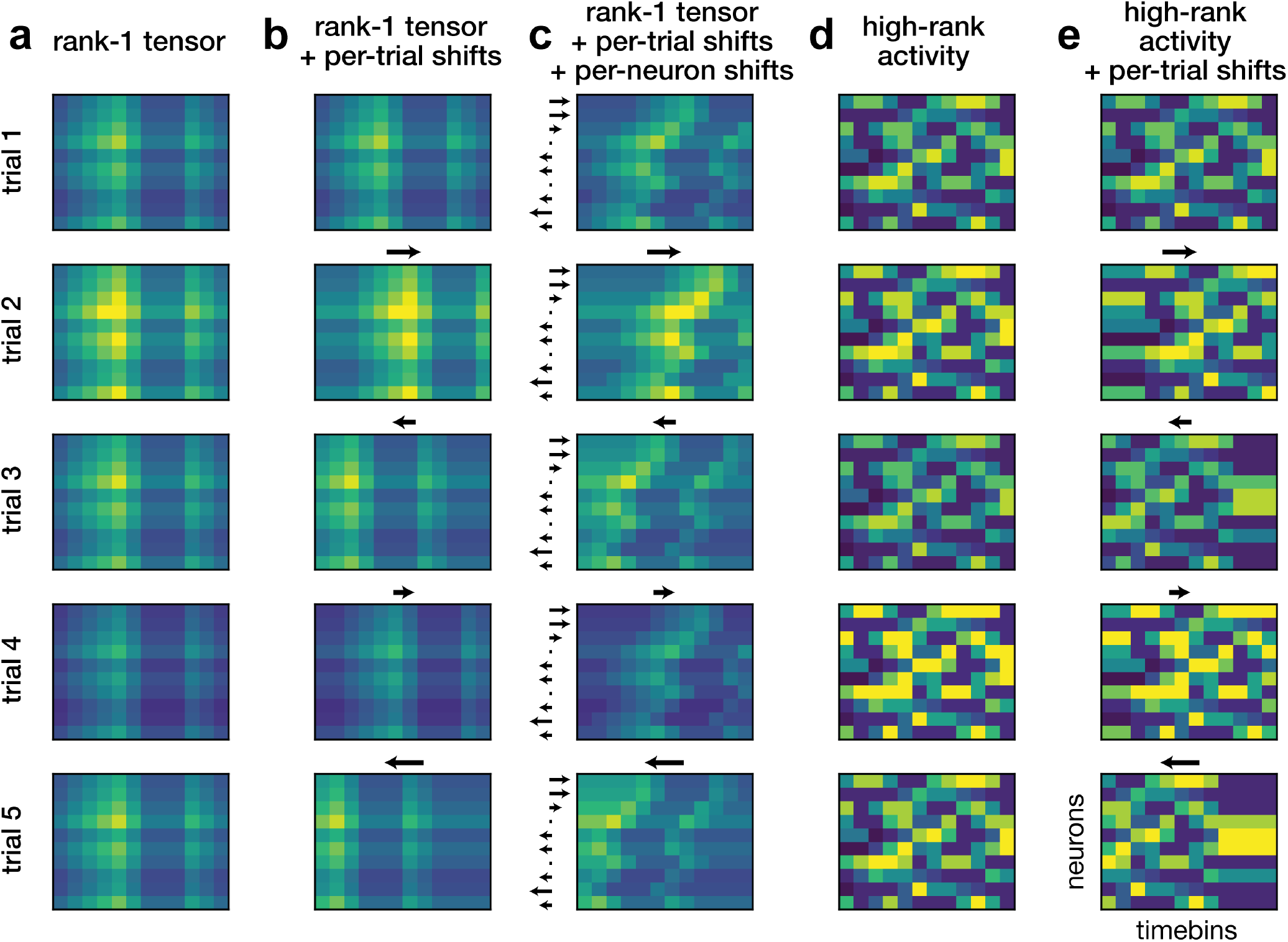
Examples of population dynamics and single-trial variability. Each panel shows five trials of population activity as a neurons *x* timebins heatmap—i.e., each heatmap visualizes an *N × T* matrix **X**_k_ of single-trial activity. (A) Activity according to a rank-1 tensor decomposition model. (B) Activity according to a rank-1 tensor decomposition model with per-trial shift parameters (equivalent to eq. (8) with *α_nr_ =* 0). The activity on each trial is still a rank-1 matrix, but the resulting tensor contains higher-dimensional structure across trials. Black arrows above each trial denote the direction and magnitude of the shift relative to trial 1. (C) Activity according to a rank-1 tensor decomposition model with per-neuron and per-trial shift parameters (equivalent to eq. (8)). The tensor is now high-dimensional across neurons as well as trials, despite being related to panel A by a simple transformation. Arrows above each trial denote per-trial shifts as in panel B. The direction and magnitude of per-neuron shifts are denoted by the arrows to the left of each neuron. (D) High-dimensional neural activity that is repeated over trials; the only difference across trials is an overall amplitude/scale parameter. Though the across-trial variability is low-dimensional, any tensor decomposition would have trouble fitting these data due to the complex intra-trial dynamics. (E) Same as panel *D*, but with per-trial time shifts. Note that trial-averaging in this setting would produce an inappropriate estimate of the single-trial dynamics; a shift-only time warping model (eq. (4)) can correct for the temporal variability across trials and produce a better estimate.

Other datasets may be less “close” to a low-rank tensor model. Figure 3d illustrates an example dataset where neural dynamics within each trial are high-dimensional, but are nonetheless reliably repeated across trials. Such data could be generated, for example, by visual cortex in response to movies of natural scenes (Stringer et al. 2019). In this case, the within-trial dynamics are not easily compressed by tensor decomposition or any other method. However, in the presence of noise, one can still average across trials to get a reliable estimate of these high-dimensional dynamics. Figure 3e illustrates the same dataset as panel *d*, but with the addition of random delays on each trial. Here, trial-averaging will fail to produce an accurate estimate of the within-trial neural dynamics due to misalignments across trials. A time warping model, which we describe next in section 1.3, aims to explicitly model and overcome these misalignments and may be applicable even when within-trial dynamics are high-dimensional.

#### 1.2.1 A note on nonnegative factorizations

Nonnegative tensor decomposition (Lim and Comon 2009; Paatero 1997) is a higher-order analogue of the well-known nonnegative matrix factorization model (Lee and Seung 1999). The structure of this model is identical to eq. (2), but the neural factors (*u_nr_*), trial factors (*v_kr_*), and temporal factors (*w_tr_*) are all constrained to be greater than or equal to zero during optimization. In Williams et al. (2018), we reported that this constraint improves the consistency and interpretability of tensor decomposition without greatly hurting the model’s accuracy. Section 2 describes optimization methods for enforcing nonnegativity constraints in the context of tensor decomposition and time warping models. Incorporating nonnegativity has been useful in the vast majority of practical applications that I’ve encountered. Thus, one may safely read this manuscript with the assumption that *u_nr_* ≥ 0, *v_kr_* ≥ 0, and *w_tr_* ≥ 0 for all *n, t, k*, and *r*, as this case is often the rule, rather than the exception.

### 1.3 Time Warping

To capture variability in timing across trials, Williams et al. (2020) proposed the following **time warping model**:

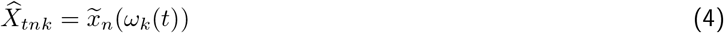

Here *ω_k_(t)* is a monotonically increasing **warping function** mapping integers {1, 2,…, *T*} to real numbers on the interval [1,*T*]. The vector 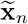 is a length-*T* vector holding the *response template* for each neuron *n*. The response templates for each neuron, 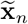, and the warping functions for each trial, *ω_k_(t)*, are optimized to minimize the least-squares criterion in eq. (1). The response templates are fit independently for each neuron, and thus high-dimensional within-trial dynamics can be accommodated by this model (see fig. 3e).

Since *ω_k_(t)* is not generally an integer it cannot be used as an index into the discrete vector 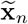. Equation (4) thus contains an implied **linear interpolation** step:

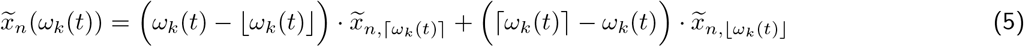

where 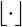 and 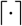 respectively denote flooring (rounding down to the nearest integer) and ceiling (rounding up to the nearest integer) operations. We will continue to use parenthetical indexing—e.g., *x(t)*—to denote indexing with linear interpolation, and subscript indexing—e.g., *x_t_*—to denote direct indexing with integer variables.

By its virtue of being linear, the interpolation step in eq. (5) can be expressed as a matrix multiplication. Thus, every warping function, *ω_k_(t)*, is uniquely associated with a warping matrix, Ω_*k*_. For example, suppose the warping function on each trial has the form *ω_k_(t)* = *t* + *β_k_*, where *β_k_* is a per-trial shift parameter. In Williams et al. (2020) we refer to this a **shift-only warping model**. Of course, when *β_k_* = 0, we have *ω_k_(t)* = *t* and Ω*_k_* is a *T* × *T* identity matrix. The warping matrices associated with some other example warping functions are show below:

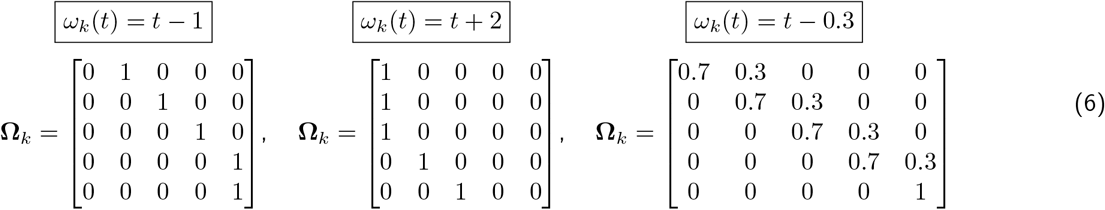

Using this notation, we can reformulate eq. (4) as follows:

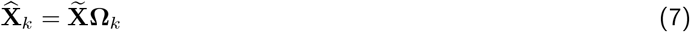

which is the time warping analogue to eq. (3). Note that the above shift model clamps the endpoint values of the response template; if desired, this can be modified to accommodate periodic boundary conditions (Mørup et al. 2008).

In contrast to tensor decomposition, the time warping model defined in eq. (4) does *not* assume that firing rate dynamics are low-dimensional, since a different firing rate template is learned for each neuron. Further, by transforming the time axis on each trial by a learned warping function, the model can account for variability in the onset and duration of neural dynamics.

On the other hand, there are some drawbacks to this time warping model. The model does not explicitly account for trial-by-trial variations in amplitude, which, as discussed in section 1.2, are often of interest. Perhaps more importantly, the time warping model used by Williams et al. (2020) assumes that *all neurons share the same warping function on a trial-by-trial basis* and thus does not explore potential variations in timing expressed across multiple sub-populations of neurons. Thus, the time warping and tensor decomposition models have complementary strengths and weaknesses. The following two sections discuss modeling extensions that aim to achieve the best of both models.

### 1.4 Time-Shifted Tensor Decomposition

As discussed in section 1.2, there are two principal “failure modes” in which tensor decomposition *nearly* works, but fails due to temporal misalignments. First, neurons may fire with slightly different latencies to each other, for example in a short temporal sequence (see Mackevicius et al. 2019 and references therein). Second, the ensemble itself may fire at a different latency on a trial-by-trial basis. These possibilities were illustrated in fig. 3b-c.

One way to correct these discrepancies is by incorporating additional time warping functions into each component of the tensor decomposition. The resulting model can be very complex if each time warping function is allowed to be highly nonlinear; however, a key empirical observation in Williams et al. (2018) is that very simple forms of time warping are sufficient to uncover interesting features in neural data. Thus, we will introduce the simplest form of time warping, *shift-only warping*, leading to the following **time-shifted tensor decomposition** model (see Harshman et al. 2003; Mørup et al. 2008 for related prior work):

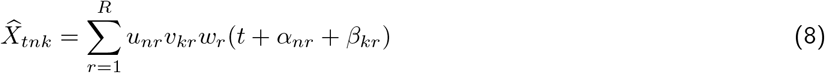

Here, the parenthetical indexing notation, *w_r_*(·), denotes linear interpolation as in eq. (5). In addition to optimizing the low-dimensional factors {*u_nr_*, *v_kr_, w_tr_*}, we now must also optimize *α_nr_* and *β_kr_* which may be interpreted as a per-neuron and per-trial shift parameters for each low-dimensional component. The time-shifted tensor decomposition has a total of (2NR + *KR + 2TR)* parameters, which is a very modest increase over the number of parameters in vanilla tensor decomposition.

Equation (8) can be reformulated as:

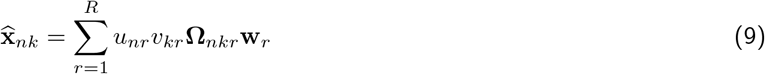

Where **Ω***_nkr_* denotes a time warping matrix as in eq. (6). Note that a different warping function is applied to each low-dimensional temporal factor, **w**_r_, for every trial and neuron.

Optimizing time-shifted tensor decomposition is potentially much more challenging than vanilla tensor decomposition, due to the additional shift parameters which are not easily fit by gradient-based methods.^1^ Nonetheless, we have found this to be possible in practice, as long as the magnitudes of the shift parameters are not too large.

### 1.5 Multi-Shift Model

The time-shifted tensor decomposition still assumes that the same temporal firing rate function (i.e., the temporal factors) are shared across all neurons in an ensemble, up to a temporal shift. This assumption may be too restrictive for neural dynamics that span many dimensions (Stringer et al. 2019). We can relax this assumption by replacing the low-dimensional neuron and temporal factors used in tensor decomposition (*u_nr_* and *w_tr_*), with firing rate templates, as used in the vanilla time warping model 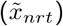. This results in the **multi-shift model**:

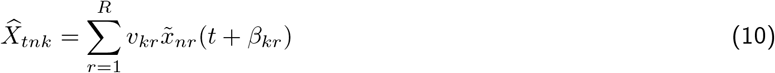

Using the warping matrix notation, the model can also be viewed as:

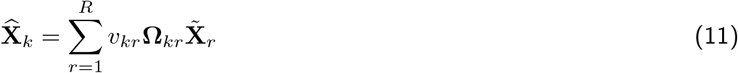

This model extends the power of a vanilla shift-only time warping model by introducing multiple cell ensembles (indexed by *r*), which are independently modulated both in amplitude (by trial factors, *v_kr_*) and in timing (by per-trial shift parameters *β_kr_*). The multi-shift model contains *RNT* + *2KR* parameters, which can be considerably larger than time-shifted tensor decomposition in large-scale neural recordings. Nonetheless, since the number of total parameters grows slowly as *K* increases, the model is still often feasible to work with in practical circumstances. Additionally, we can enforce nonnegativity constraints on 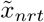 in a manner analogous to tensor decompositions (see section 1.2.1); this mitigates the potential for overfitting by limiting the expressivity of the model.

In summary, we have outlined two unsupervised learning models to model single-trial data. The first (eq. 8) closely resembles tensor decomposition, and incorporates shift-only time warping across neural and trial dimensions. The second (eq. 10) more closely resembles a time warping model, but incorporates multiple dynamical templates with per-trial amplitude modulation.

### 1.6 Choosing the number of components

Choosing the number of model components, *R*, is a challenging and well-known problem. In practice, this problem is often ill-posed in the sense that that there is rarely a “true” value of *R*, since our models are almost always *misspecified—*i.e., the data generation process does not follow the exact structure of our proposed model.

Nonetheless, there are a couple simple diagnostic tools to help practitioners decide on a reasonable guess for R. First, one can plot the model error as a function of *R*, and visually identify an inflection or “knee” in this monotonically decreasing curve (this visualization is called a *scree plot* in the context of PCA). A more rigorous approach is to use cross-validation: the data is split into separate training and testing partitions, which are respectively used for parameter fitting and model comparison. In this case, the model test error should eventually begin to increase for large enough values of *R*, indicating overfitting.

Second, one can use so-called *model stability* measures to choose an appropriate model. In this context, a model is considered “stable” if it consistently converges to the same solution across multiple optimization runs from different random initializations. Conversely, a model is “unstable” if it converges to different solutions. Stability can be quantified by quantifying the similarity (e.g. with correlation coefficients) across optimization runs.

Stability criteria have previously been used to choose the number of components in clustering models (Luxburg 2010) and matrix factorization models (S. Wu et al. 2016). The presumption of this approach is that over-parameterized models (i.e. those with *R* too large) are less stable than well-tuned models. Intuitively, this occurs because models with excess degrees of freedom can identify degenerate, redundant solutions, while models with fewer parameters are more constrained. Indeed, under weak assumptions the global solution of tensor decomposition is “essentially unique” (ignoring permutations and rescalings of factors) for small enough values of *R*; this uniqueness is eventually lost as *R* increases (Kruskal 1977; Rhodes 2010).^2^ In the context of exploratory data analysis, model stability is a clearly desirable feature—if multiple solutions with similar approximation error exist, then it becomes challenging to interpret and bestow meaning onto any particular solution.

Given two tensor decomposition models, with parameters {**u**_*r*_, **v**_*r*_, **w**_*r*_} and 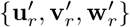, Williams et al. (2018) used the following similarity metric to quantify stability:

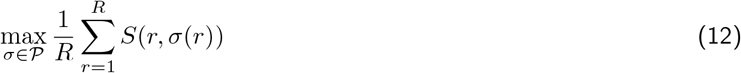

where 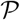 is the set of all permutations of length *R*, σ(·) denotes one such permutation, and *S(i,j)* computes the similarity between component *i* in model 1 and component *j* in model 2 as follows:

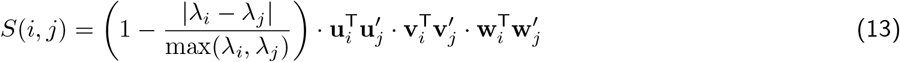

here *λ_r_* is a scale representing the norm of the *r*^th^ component found by normalizing all factors **u**_*r*_, **v**_*r*_, **w**_*r*_ to unit length; see T. Kolda and Bader (2009). This similarity measure can be efficiently computed by enumerating an *R* × *R* matrix of pairwise similarity scores between all pairs of factors; then the optimal permutation of the factors, σ, can be found by the Hungarian algorithm in *O*(*R*^3^) running time (Jonker and Volgenant 1987; Burkard et al. 2012).

Equation (13) must be modified to be applicable to time-shifted tensor decomposition models. In particular, it is difficult to directly compare the temporal factors between two models due to invariances in the model structure. For example, adding a constant to the per-trial shift parameters, e.g. *β_kr_* ← *β_kr_* + 1 for all *k*, could result in very little change in the model prediction if the values in **w**_r_ are shifted in the opposite direction, i.e. *w_r_(t)* ← *w_r_(t + 1)* for all *t*.^3^ To circumvent this problem we can compute similarity by defining:

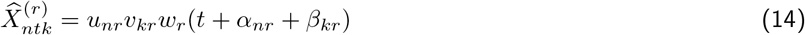

as the contribution of component *r* to the model’s reconstruction. Then, we define the similarity between component *i* in model 1 and component *j* in model 2 as the norm of the residuals, 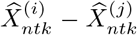, where the first term is computed using the first model’s parameters {**u**_*i*_, **v**_*i*_, **W**_*i*_,α_*ni*_, β_*ki*_} and the second term is computed using the second model’s parameters 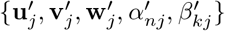. Using this new definition of 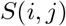, which accounts for invariances introduced by the shift parameters, the optimal permutation can be computed as described above and eq. (12) can be applied to compute an overall similarity score between the two models.

## 2 Parameter Estimation

This section describes optimization methods to fit the four models described in section 1. Readers focused on higher-level concepts may wish to skim or skip this section entirely, as it contains somewhat exhaustive, step-by-step derivations of the parameter update rules for those interested in deeper technical details. Python implementations of these parameter estimation routines are provided at: https://github.com/ahwillia/tensortools

For a quadratic loss function, we can derive efficient and exact coordinate-wise updates for all model parameters, except for the shift parameters involved in time warping. These coordinate descent routines are known as hierarchical alternating least squares (HALS) in the matrix and tensor factorization literature (Cichocki et al. 2007; Gillis and Glineur 2012). The key idea is to focus on updating one component at a time, which reduces the complex, nonconvex optimization problem to solving a series of easy-to-solve problems. Additionally, optimizing over single components makes it very simple to enforce nonnegativity constraints on parameter values.

### 2.1 Tensor Decomposition

In the case of a vanilla tensor decomposition (starting from eq. (2)) the objective function is:

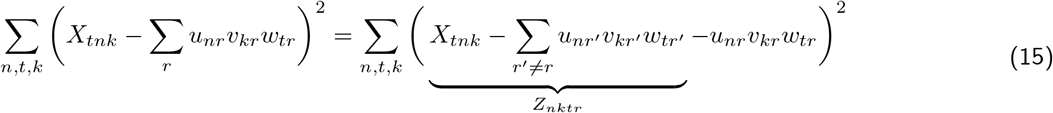

where we have defined *Z_nktr_* as the residual, excluding the r^th^ component of the model. If we are optimizing over any parameter in the r^th^ component, then the *Z_nktr_* term is a constant. Now we can focus on minimizing:

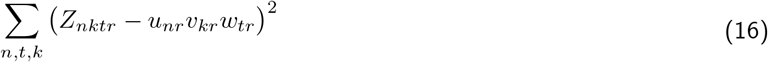

which can be done in closed form. For example, consider optimizing over *w_tr_* while treating the other factors (*u_nr_* and *v_kr_*) and other components (all parameters where *r’* ≠ *r*) as fixed constants. The objective function is convex in *w_tr_*, so there is a unique minimum, at which the gradient must be zero. A short calculation reveals this optimal value for *w_tr_*:

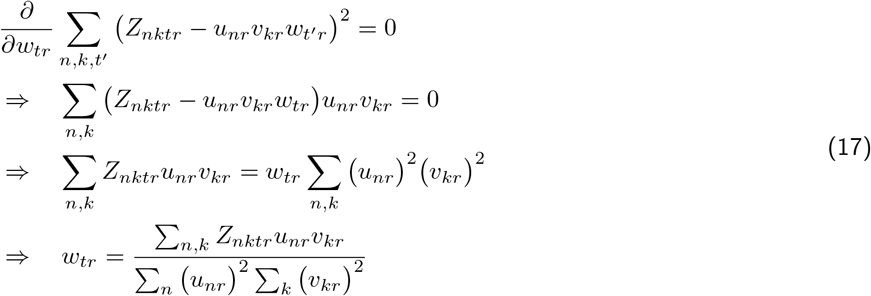

Further manipulation of the numerator reveals that we need not form the residual tensor, *Z_nktr_*, explicitly since:

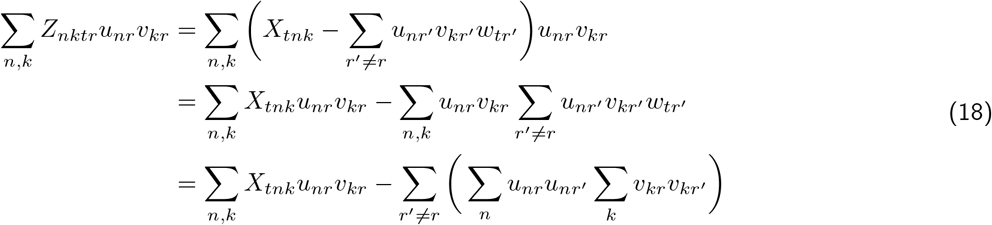

It is worthwhile to connect these derivations with the terminology and notation in T. Kolda and Bader (2009) and related works. For example, the term 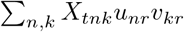 represents a matrix multiplication between an unfolded (matricized) tensor and a Khatri-Rao product of vectors **u**_r_ and **v**_r_. To keep a streamlined narrative, we will not cover these connections in any further detail.

The tensor decomposition model we consider has a clear symmetry across the three sets of low-dimensional factors. Thus, we can easily repeat the above derivation for the other factors, *u_nr_* and *v_kr_*, and arrive at the following update rules:

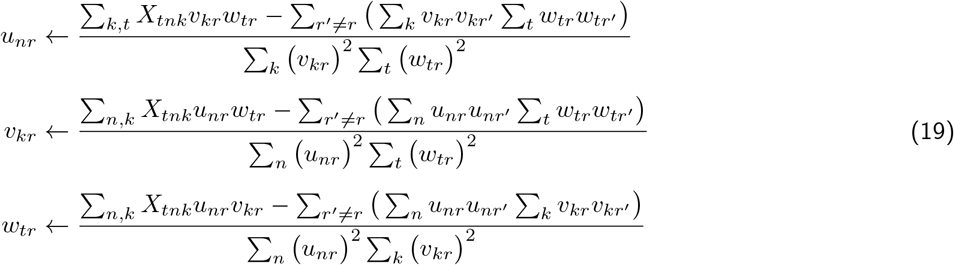

In practice, we often incorporate **nonnegativity constraints** on the low-dimensional factors for the purposes of interpretability and identifiability. By appealing to the Karush–Kuhn–Tucker (KKT) conditions, one can show that incorporating these constraints simply involves truncating any negative values to zero:

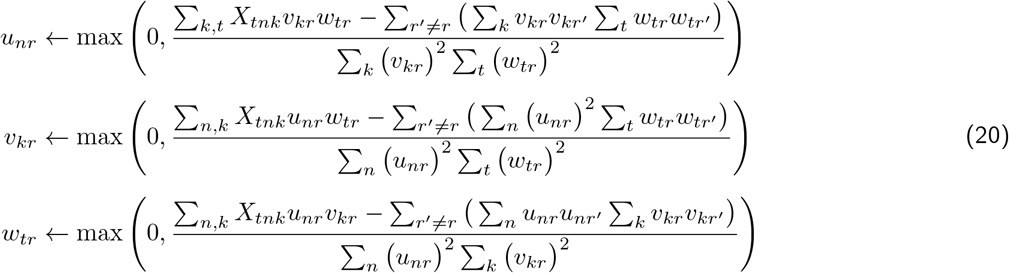

It should be emphasized that these updates are **not** valid when multiple components are optimized at once—in this case, the updates can no longer be written in closed form and involve solving a quadratic program (nonnegative least-squares) problem (Gillis 2014; J. Kim et al. 2014).

### 2.2 Time Warping

Utilizing the warping matrix notation (see eq. 6), we can formulate the time warping model’s objective function as:

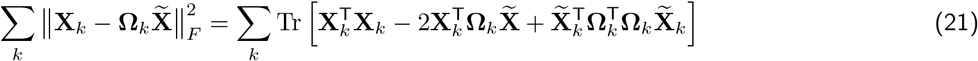

We adopt a block-coordinate descent approach, first treating the warping matrices, **Ω***_k_*, as fixed and optimizing 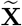. As before, we can find a closed form update by identifying where the gradient is zero:

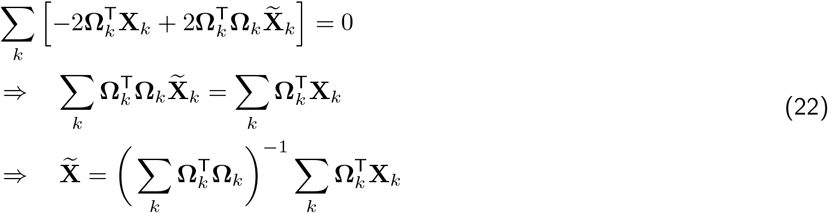

This update is very efficient to compute, as it can be shown that each 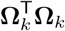 is symmetric and tridiagonal (see Williams et al. 2020 for details). In python, such linear systems can be efficiently solved by the `scipy.linalg.solveh_banded` function. Note that there is no reason to enforce nonnegativity on the warping template **X** as it will naturally be nonnegative for nonnegative data.

After updating the response templates according to eq. (22) we must then update the per-trial warping functions. Unfortunately, this can be quite complicated. The optimal *nonlinear warping* can be found by the celebrated *Dynamic Time Warping* algorithm (Berndt and Clifford 1994). Such nonlinear warping paths can be very complex and are thus susceptible to overfit to noise, prompting ongoing work on regularized and smoothed time warping functions (Cuturi and Blondel 2017; Duncker and Sahani 2018).

In Williams et al. (2020), we constrained the warping functions to be piecewise linear. Empirically, this simple class of functions performed well on many neural datasets. Furthermore, these functions are specified by a small number of learned parameters—e.g. a piecewise linear function with two segments contains four free parameters. In practice, shift-only warping or linear warping (containing one and two free parameters, respectively) often perform quite well.

Optimizing the warping functions is simple when we restrict them to be piecewise linear. One could use gradient descent, which would involve computing derivatives of the model loss with respect to the knots defining the piecewise linear segments. However, when neural firing rates are oscillatory, this loss landscape can exhibit suboptimal local minima. Instead, we optimized by a brute force randomized search (Bergstra and Bengio 2012) over this low-dimensional parameter space. Specifically, the piecewise linear functions are randomly perturbed and the parameters are updated whenever this perturbation improves the model loss. Further details are provided in Williams et al. (2020).

### 2.3 Time-Shifted Tensor Decomposition

Utilizing warping matrices, we can formulate the model’s objective function as:

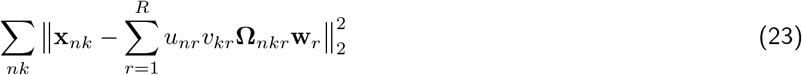

We follow a similar strategies to those outlined in sections 2.1 and 2.2. First, to optimize the warping functions, we perform a randomized searches as described in section 2.2. Recall from eqs. (6) and (8) that we assume the warping functions to correspond to per-neuron shifts, *α_nr_*, and per-trial shifts, *β_kr_*. Due to interactions between these two sets of parameters, it is not easy to simultaneously optimize *α_nr_* and *β_kr_* in parallel. However, we can perform update these parameters in two blocks—first over neurons, then over trials—each done parallel.

To optimize the remaining parameters (the low-rank neural, temporal and trial trial factors), we follow the approach of section 2.1 and consider optimizing a single component at a time. Denote the residual for neuron *n* on trial *k* as 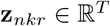; that is:

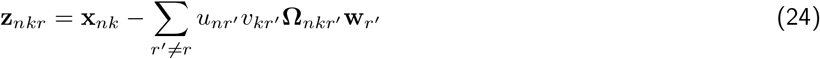

The objective function with respect to the r^th^ component can be expressed as:

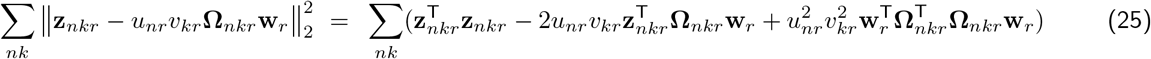

As before, closed form parameter updates can be derived for the neural, temporal, and trial factor by identifying where the gradient is zero.

For the temporal factor, **w**_*r*_, we obtain:

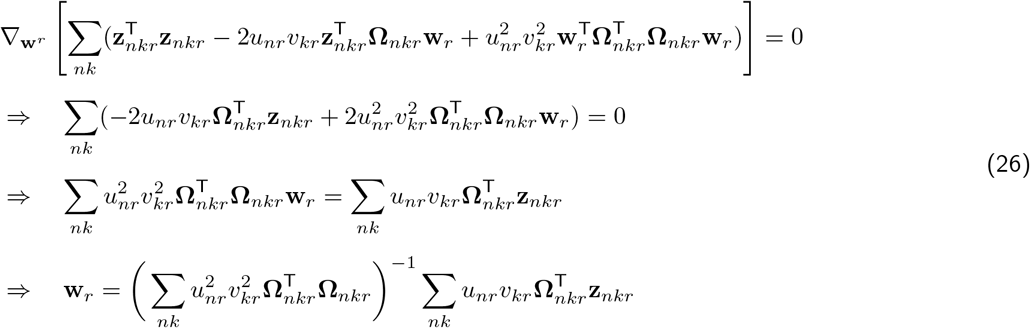

which can be viewed as a re-weighted version of eq. (22).

For the trial factor, **v**_*r*_, we obtain:

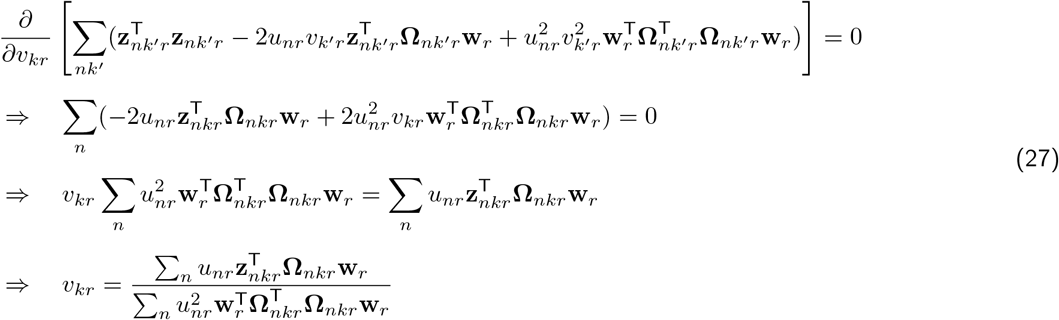

Deriving an analogous update rule for the neuron factors follows a nearly identical series of computations, ultimately obtaining:

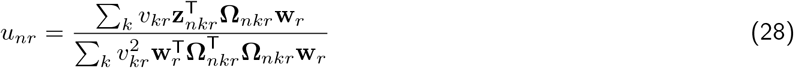

In practice, we find that it is beneficial to constrain the neural and trial factors to be nonnegative. Enforcing this constraint is simple and can be done in analogy to eq. (20). Enforcing nonnegativity on the temporal factors is a bit more challenging, since the corresponding update rule (derived in eq. 26) involves multiple variables at once. As discussed in Gillis (2014) and J. Kim et al. (2014), a more appropriate update rule must use *nonnegative least squares* methods. This could be accomplished by projected gradient descent, as we’ll see in the following section (see eq. (34)). Alternatively, the nonnegativity constraint could be ignored on the temporal factors—in practice, constraining the neural and trial factors to be nonnegative may be sufficient to produce a stable and interpretable model.

### 2.4 Multi-Shift Model

Optimizing the multi-shift model can also be accomplished by coordinate descent approach. First, define the residual matrix 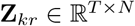 as:

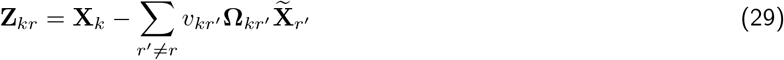

for trial *k*. Then, to update component *r*, we can focus on minimizing the objective function:

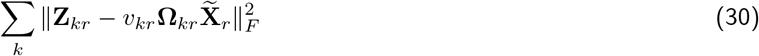

Again, we resort to a randomized search over per-trial shift parameters to optimize **Ω***_kr_*. We have found it useful to constrain both the trial factors, *v_kr_*, and response templates, 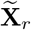, to be nonnegative. Updating *v_kr_* under this constraint can be done in closed form, but we resort to a fast projected gradient descent routine to update the response templates.

The update rule for *v_kr_* is found by identifying where the gradient is zero:

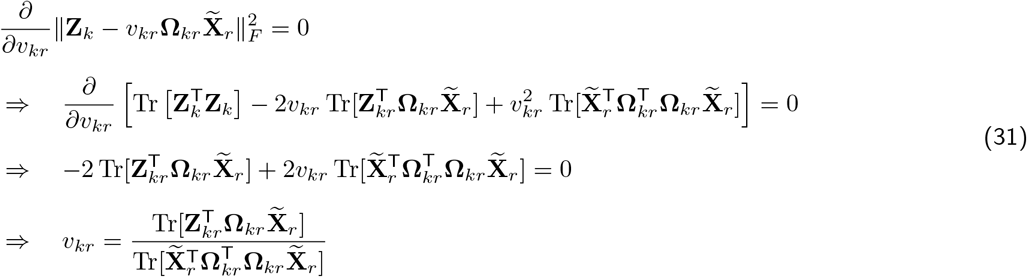

Since we are optimizing over a single variable, it is valid to simply truncate negative values to enforce *v_kr_* ≥ 0. In summary, the update rule, which can be computed in parallel across trials, is:

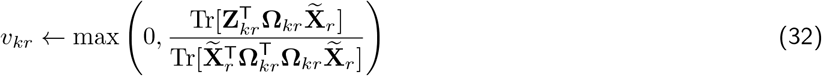

Finally, we must derive an update rule for the response templates, 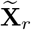. We begin by deriving the gradient:

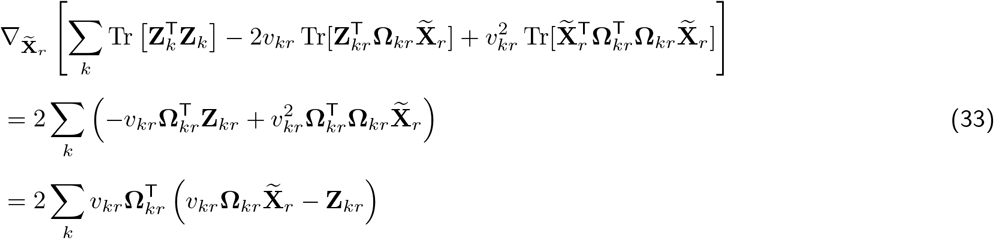

By differentiating once more can find the Hessian matrix with respect to 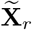. The Hessian can be viewed as a *NT × NT* block diagonal matrix with 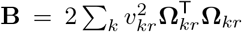 repeated *N* times as the blocks.^4^ Since the objective function is quadratic, the gradient is globally Lipschitz continuous with constant *L* = 2||**B**||_2_, with ||·||_2_ denotingthe largest eigenvalue of a matrix (operator norm). Standard convex optimization theory tells us that projected gradient descent will converge as long as the step size is less than this Lipschitz constant (Boyd and Vandenberghe 2004). We can compute a very tight upper bound on this maximum eigenvalue very cheaply via the Gershgorin circle theorem. Recall that **B** is tridiagonal, nonnegative, and symmetric, due to the structure of the warping matrices. Thus, the maximum eigenvalue must be a real number and be less than *γ* = max(diag(**B**))) + 2max(offdiag(**B**)). We set the stepsize of gradient descent to be the inverse of this upper bound, assuring fast convergence.^5^ In summary, we update 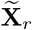 according to:

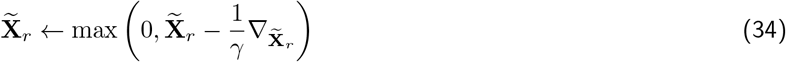

with the gradient term computed according to eq. (33).

## 3 Demonstrations on Synthetic Data

As mentioned at the beginning of section 2.2, tensor decomposition can fail to uncover the intended low-dimensional structure of data when the timing of neural dynamics is variable. Figure 4 demonstrates this in a simulated dataset with *N* = 60 neurons, *T* = 120 timebins, and *K* = 200 trials. Poisson-distributed binned spike counts were simulated according to the 2-component ground truth model shown in Figure 4a, plus random, per-trial time-shifts ±15% of the full trial duration. The maximum Poisson rate parameter in any timebin was less than 0.3, so that the simulated spike patterns were highly sparse and divergent from the Gaussian likelihood/quadratic loss criterion associated with the models (see eq. (1)). The minimum Poisson rate parameter in any timebin was 0.02, so that some additional spikes were present as “background noise.”

**Figure 4:**
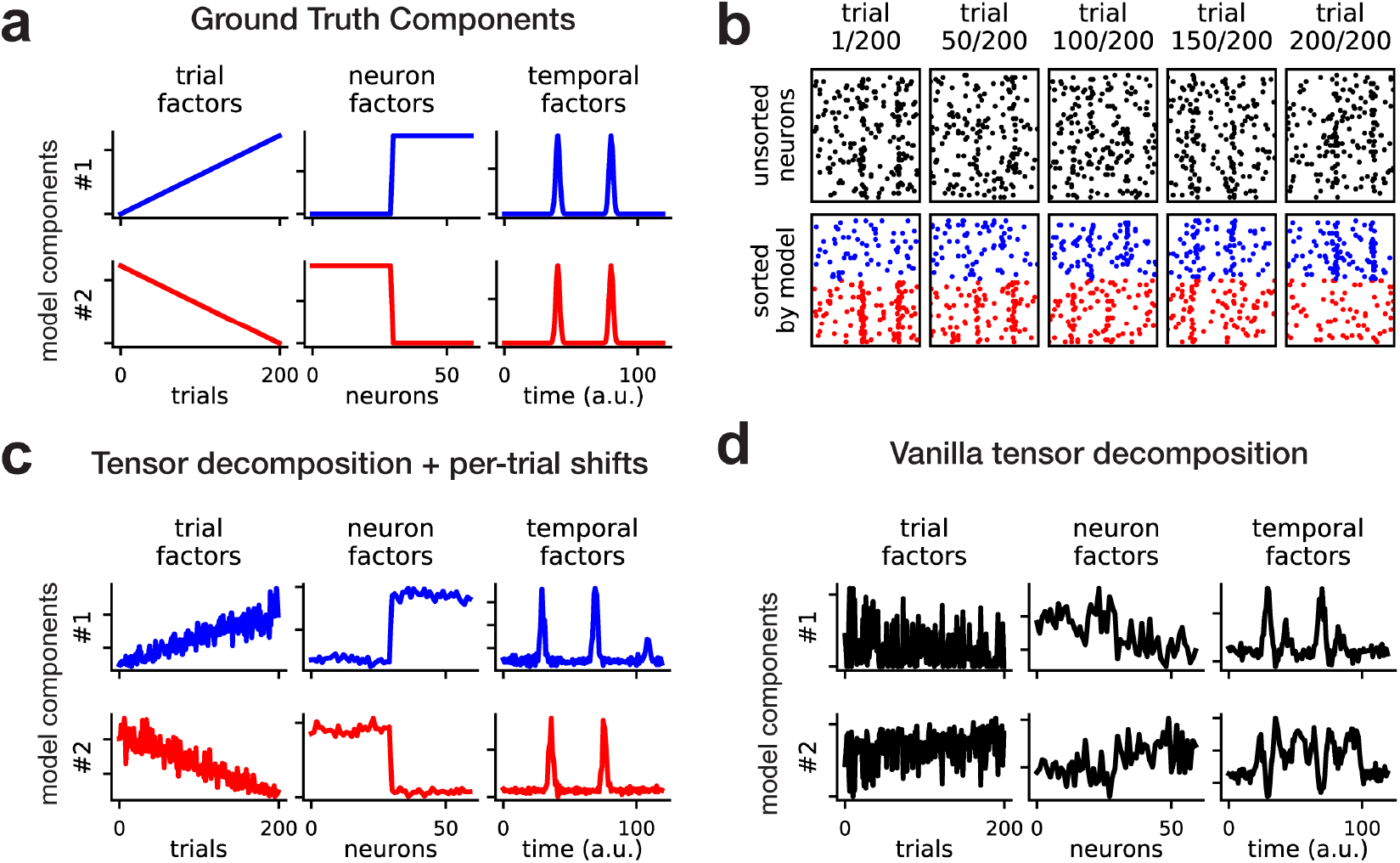
Synthetic spiking activity of two neural populations with random per-trial latencies. The ground truth spiking probabilities follow eq. (8) with *α_nr_ =* 0 for each neuron *n* and component *r*, and *β_kr_* randomized ±15% of the trial duration for each trial *k* and component *r*. (A) Ground truth components for simulated data. The *β_kr_* delay parameters are not visualized. Component 1 and 2 are colored blue and red, respectively. (B) Simulated spike data of *N* = 60 neurons over *K* = 200 trials. Spike counts were sampled from a Poisson distribution, with time-varying firing rate function given by the model in panel A. Top row (black spikes) shows trials 1, 50, 100, 150, 200 with neurons shown in a randomized order. Bottom row (blue and red spikes) shows the same data with neurons grouped according to the learned neuron factors. The first component (in blue) grows in amplitude over the course of the simulated session, while the second component (in red) decreases in amplitude over the course of the session. (C) Low-dimensional factors recovered by the shifted tensor decomposition model (the learned *β_kr_* parameters are not visualized). The factors are colored to match panels A. (D) A tensor decomposition model without time warping fails to recover the ground-truth structure of the model. The factors do not match panel A and thus are left uncolored.

The factors in Figure 4a can be interpreted as follows. First, as indicated by the neuron factors (middle column), there are two non-overlapping sub-populations or ensembles of cells. Second, as indicated by the temporal factors (right column), each ensemble fires twice within every trial—the responses are sharp and only last for a short time interval, making them somewhat difficult to detect, especially in the presence of trial-to-trial variation in response onset. Finally, as indicated by the trial factors (left column), the first neural ensemble grows linearly in amplitude over trials, while the second neural ensemble diminishes linearly over trials.

Figure 4b shows the activity of all neurons in five trials equally spaced across the full session of *K* = 200 trials. In the top row of raster plots, each of the *N* = 60 neurons are assigned to a random vertical coordinate to simulate the appearance of raw experimental data. It is very hard to visually identify recurring temporal patterns within trials and longer-term changes in population activity across trials. The bottom row of raster plots shows the same data, but with neurons re-sorted and colored by the low-dimensional neuron factors identified by time-shifted tensor decomposition in Figure 4c. The model successfully groups neurons into their two ground truth ensembles, enabling one to visually identify the double banded structure in each ensemble’s response pattern and the fact that ensemble #1 (in blue) intensifies over the session while ensemble #2 (in red) decreases in amplitude. The trial-to-trial variability in response timing is also visible upon close inspection of these re-organized raster plots. Overall, the time-shifted tensor decomposition model captures the correct qualitative structure of the simulated dataset (compare fig. 4a and fig. 4c). In contrast, a classic tensor decomposition model (see eq. (2)) fails to identify any of this interpretable structure in the neural data (fig. 4d); this is due entirely to the variability in neural response times.

Next, we performed a hyperparameter sweep to determine whether the diagnostic tools described in section 1.6 could successfully identify the number of components and the scale of the per-trial shifts in a similar synthetic dataset.^6^ We fit models with *R* ∈ {1, 2,3,4, 5}. Further, we set a maximum absolute value to the per-trial shift parameters, *β_kr_*; expressed as a fraction of the total trial duration, these limits were *β_max_* ∈ {0,0.04,0.08,0.12,0.16, 0.2}. Note that *β_max_* = 0 corresponds to traditional tensor decomposition. For each combination of *R* and *β_max_* we fit seven models from different random initializations. Figure 5a shows that a model with *R* = 2 components (yellow line) with *β_max_* = 0.16 closely matches the lowest error achieved by any model; the hyperparameters of this model were the closest to the ground truth (which has *R* = 2 and *β_max_* = 0.15). Furthermore, when *β_max_* was set lower than its ground truth value, even models with additional components (*R* > 2) often performed suboptimally. Note, however, that for traditional tensor decomposition (i.e., *β_max_* = 0) the model with *R* = 5 components noticeably outperforms models with fewer components. Thus, the presence of per-trial temporal shifts can cause us to mistakenly believe that high-rank tensor model is needed; introducing shift parameters into the tensor decomposition model reveals that *R* = 2 components is necessary, resulting in a simpler and much more interpretable model with similar (or even superior) performance.

**Figure 5:**
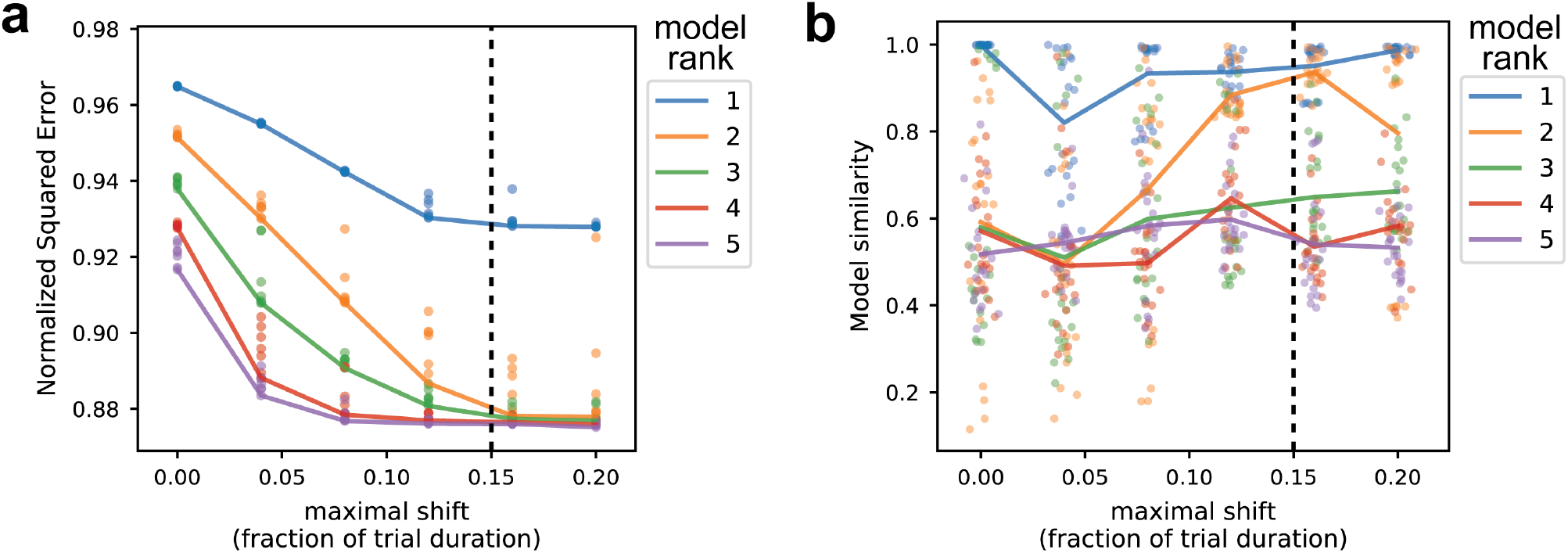
Hyperparameter sweep for time-shifted tensor decomposition. Colored lines denote models with different numbers of components (i.e., *R*, the rank of the tensor decomposition), while the horizontal axis measures the maximal per-trial shift, *β*_max_. The ground truth model had *R* = 2 components (yellow line) and *β*_max_ = 0.15 (vertical dashed black line). For simplicity, no per-neuron shifts (*α_nr_* in eq. (8)) were included in the model. (A) Normalized reconstruction error, 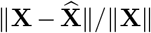, of all models. (B) Model similarity scores (see section 1.6) across different random initializations of the same hyperparameter set.

Figure 5b shows the model stability criterion (see section 1.6) over the explored hyperparameter range. Every dot in fig. 5b corresponds to the similarity score (ranging between zero and one) between a pair of models with the same hyperparameters. Models with *R* = 1 components were the most stable; however, these models had suboptimal performance as already shown in fig. 5a. Models with too many components (*R* > 2) were consistently unstable. Interestingly, models with the correct number of components (*R* = 2) was not always stable—if the maximal per-trial shift parameter was less than the ground truth value, the similarity between model fits was consistently low. The model closest to the ground truth (*R* = 2 and *β_max_* = 0.16) demonstrated relatively high stability, which, in conjunction with the results in fig. 5a, provides strong evidence for its superiority.

## 4 Conclusion and Future Work

Taken individually, tensor decomposition and time warping can be broadly applicable and insightful methods for neural data analysis (Williams et al. 2018; Williams et al. 2020). Nonetheless, there may be cases where these two models are insufficient. Neural ensemble activity may be shifted across neurons or on a trial-by-trial basis, hampering the interpretability of tensor decomposition. On the other hand, the time warping model described in Williams et al. (2020) assumes that all neurons shared the same trial-by-trial warping function and have a single canonical response pattern. We described how to combine the complementary strengths of these two models into two new models: time-shifted tensor decomposition, and a multi-shift model. Previous work has explored similar modeling ideas, but with different proposed algorithms and motivating applications (Harshman et al. 2003; Mørup et al. 2008). The unified treatment of these hybrid models provided here should clarify the conceptual connections between these previous works. Additionally, we provided an open source Python implementation for these shifted tensor decomposition models at: https://github.com/ahwillia/tensortools

## Acknowledgements

I wish to thank Tammy Kolda (Sandia National Labs) for introducing me to tensor decomposition methods and Surya Ganguli (Stanford) for helping me relate these ideas to a neuroscience audience. Jordan Sorokin (Stanford) helped me develop and beta-test early versions of the time-shifted tensor decomposition code. Jordan Sorokin, John Huguenard, and Isabel Low (all Stanford) provided experimental data and feedback that helped me develop these models—though no experimental data is discussed in these notes, their input nonetheless helped form and motivate the data analysis approaches described here. Scott Linderman and Jimmy Smith (both Stanford) provided feedback on the manuscript. I also wish to thank the U.S. Department of Energy CSGF program for supporting my PhD research on these topics.

## Footnotes

1 While gradients can be computed for these parameters, the model may exhibit suboptimal local minima when the temporal factors are multimodal.

2 Note that while the global solution is provably “essentially unique” for small enough *R*, we can only guarantee that iterative optimization methods converge to a local minimum. Thus, even when the uniqueness conditions of Kruskal (1977) are met, tensor decomposition may still exhibit some instability due to convergence to distinct local minima. Nonetheless, in practice, instability tends to increase as *R* increases, and can thus be used as a heuristic to choose the number of components.

3 For the sake of intuition, we ignore the effect of boundary conditions where *t* =1 and *t = T*. These boundary effects will only create small discrepancies if *T* is large and the shifts are small.

4 In eq. (33) we treated the gradient as *T × N* matrix, and thus by common convention we would treat the Hessian as a order-4 tensor. However, if we view the gradient as a vectorized version of eq. (33), then the Hessian would be an *NT × NT* matrix as described.

5 The bound on the maximum eigenvalue is not tight and it is very easy to derive tighter approximations using the Gershgorin circle theorem. Additionally, we can use specialized eigenvalue solvers such as scìpy.linalg.eigh_tridìagonal to quickly compute the Lipschitz constant exactly. Nonetheless, the upper bound on *γ* listed here is very cheap to compute and has been sufficient in practice.

6 The signal-to-noise ratio of the simulated data in fig. 4 is near the threshold of detectability. To achieve clear results in fig. 5 we increased the width of the peaks in the temporal factors by a factor of 2. An interesting avenue of future work would be to more rigorously define the conditions (e.g. the minimal signal-to-noise ratio) under which time-shifted tensor decomposition models can be reliably fit; Kadmon and Ganguli (2018) study this problem for the classic tensor decomposition model.

